# QCAT: testing causality of variants using only summary association statistics

**DOI:** 10.1101/072355

**Authors:** Donghyung Lee, T. Bernard Bigdeli, Vladimir I. Vladimirov, Ayman H. Fanous, Silviu-Alin Bacanu

**Affiliations:** The Jackson Laboratory for Genomic Medicine, Farmington, Connecticut 06032, USA; Department of Psychiatry, Virginia Institute for Psychiatric and Behavioral Genetics, Virginia Commonwealth University, Richmond, Virginia 23298, USA; Center for Biomarker Research & Personalized Medicine; Virginia Commonwealth University, Richmond, Virginia 23298, USA; Lieber Institute for Brain Development, Johns Hopkins University, Baltimore, Maryland 21205

## Abstract

Genome-wide and, very soon, sequencing association studies, might yield multiple regions harbouring interesting association signals. Given that each region encompasses numerous variants in high linkage disequilibrium, it is not clear which are i) truly causal or ii) just reasonably close to the causal ones. Researchers proposed many methods to predict, albeit not test, the causal SNPs in a region, a process commonly denoted as fine-mapping. Unfortunately, all existing fine-mapping methods output posterior causality probabilities assuming that causal SNPs are among those already measured in the study, or have been catalogued elsewhere. However, due to technological and computational obstacles in calling many types of genetic variants, such assumption is not realistic. We propose a novel method/software, denoted as Quasi-CAausality Test (QCAT), for testing (not just predicting) the causality of any catalogued genetic variant. QCAT i) makes no assumption that causal variants are among catalogued variants, and ii) makes use of easily available summary statistics from genetic studies, e.g. variant association Z-scores, to make statistical inferences. The proposed statistical test controls the type I error at or below the desired level. Its practical application to well-known smoking association signals provide some insightful results. The QCAT software is publically available at: http://dleelab.github.io/qcat/

## INTRODUCTION

Genome-wide association studies (GWAS) and meta-analyses of GWAS have been quite successful in identifying and replicating genomic regions associated with a diverse set of human traits/diseases (1). Until recently, the National Human Genome Research Institute GWAS catalog of published GWAS has collected over 2,000 reproducibly identified association loci across hundreds of human traits/diseases (2). However, these reported GWAS loci typically harbor dozens or hundreds of statistically significant single nucleotide polymorphisms (SNPs) in high linkage disequilibrium (LD), making it fairly difficult to fine-map true causal variants out of peer non-causal ones. Consequently, among the tens of thousands reported GWAS SNPs, only very few are empirically identified as causal variants altering disease risk (3-5) and, despite considerable GWAS results, this limitation hinders the progress towards understanding biological mechanisms or pathways underlying human traits/diseases.

The first steps in the long road toward finding causal variants is the identification of larger regions of interest. Generally such a regions lie around variants harboring significant signals. However, very often signals at neighboring SNPs are correlated and, thus, they might tag a common underlying signal, i.e. they are not “independent”. To find the independent signals in a region, researchers resort to a conditional analysis in which they assess the significance of variants after conditioning on the most significant signal in the region. When genetic data is available, this conditioning can be attained by simply using the genotype at the most significant marker as a covariate and/or clumping (clustering the signals by LD) the signals (6-8). If only summary statistics are available (e.g. Z-scores), the conditioning can be achieved using only these statistics, e.g. GCTA-COJO (9). However, these are solely association based methods and, when used as fine-mapping tools, have two major disadvantages. First, they do not model at all the causality of SNPs. Due to this, if an uncatalogued causal variant with a reasonably strong signal and two of GWAS SNPs are only moderately correlated, conditional analysis might be likely to pick both GWAS SNPs as independent signals, which is obviously untrue. Second, even when the causal SNP is measured in the GWAS, it might not always be the most statistically significant.

To overcome the limitation in identifying true causal variants at GWAS-detected loci, several statistical fine-mapping methods have been proposed. For instance, a Bayesian statistical approach was used to extract “credible sets of SNPs” from GWAS association signals for 14 loci in 3 common diseases (10). The Bayesian method refines the signals by calculating the posterior probability of a SNP being causal by assuming that all causal variants are measured. While the approach is informative in prioritizing potential causal variants, it has the limitations of i) assuming only one causal variant at each loci, ii) high computational burden and ii) requiring individual-level genotype-phenotype information, which are often publicly unavailable.

To address these limitations, recently, three novel Bayesian fine-mapping methods were recently proposed: PAINTOR (11) CAVIAR (12) and CAVIARBF (13). These three methods are a big step ahead in the important conquest to fine-map association signals because they i) directly model causality in the computation of likelihood, ii) base their statistical inference only on (publicly available) marginal association statistics (e.g. two-sided Z-scores) and LD patterns (SNP genotype correlation matrices) estimated from a public reference panel, such as the 1000 Genomes data (1KG) (14). When compared to previous method, they have the advantages of: i) allowing multiple causal variants at loci, ii) faster computation time by using summary data (as opposed to subject level genetic data). CAVIAR and CAVIARBF determine credible sets of SNPs by calculating posterior probabilities of all possible subset of SNPs being casual under the assumption that all causal variants are at least known (if not already measured). Due to their greedy SNP selection procedure, their computational burden is quite demanding, rapidly increasing as the assumed number of causal SNPs at loci increases. For this reason, CAVIAR software sets the number of causal variants at most 6. PAINTOR detects potential causal variants using similar strategy used in CAVIAR and CAVIARBF, except that it provides an additional useful option to utilize functional SNP annotation information in prioritizing candidate causal SNPs. While these tools are all very useful, they all output the *relative posterior probability of causality*, i.e. under the strong simplifying assumption that all possible causal variants are measured. However, this simplifying assumption is unlikely to be met in practice (15) due to many types of genetic variation being overlooked/hard to assess even in the most favorable scenario of high coverage sequence studies (e.g. highly repetitive, hypervariable, paralogous and chromosomal duplication regions along, de novo alignment being not commonly employed etc). If an uncatalogued causal variant with a reasonably strong signal is moderately correlated with multiple GWAS SNPs, intuitively all these fine-mapping methods are likely to pick at least one of these SNPs as causal. Consequently, there is a need for methods which might output an *absolute* p-value for testing causality, which does not assume that putative causal variants are already known or catalogued.

In their present versions, all three above mentioned methods implicitly assume that the LD structures of the study cohort and the (often homogenous) reference population is identical. However, to increase sample size, and thus, the detection power, many large-scale consortia collect samples from diverse ethnic background, even including samples from different continents (16;17). Thus, for the cosmopolitan (mixed-ethnicity) cohorts, a less than careful matching of the cohort with the reference panel can results in false positive causality signals for the above three methods. Thus, the development of a method which automatically computes the LD structures for mixed ethnicity cohorts, would be welcome. Our group already developed such capability, which is already implemented in DISTMIX and JEPEGMIX software (18;19).

In this work, we propose a novel fine-mapping method. It retains all key advantages of summary statistic based fine-mapping methods by: i) allowing multiple causal variants, ii) providing high computational efficiency and iii) not requiring individual-level genotypes, while being readily applicable to mixed ethnicity cohort. We refer to our proposed method for ethnically homogeneous cohorts as **Q**uasi-**ca**usality **T**est (QCAT). (For cosmopolitan cohorts, we denote it as QCAT for MIXed ethnicity cohort (QCATMIX).) Unlike competing methods, it outputs a p-value for causality testing, which, does not assume that all possible causal variants are known or measured. We use extensive simulation studies to compare the Type I error rates and power of QCAT/QCATMIX to the fine-mapping methods, such as CAVIARBF. By applying QCAT to summary data from a large-scale meta-analysis such as the Tobacco and Genetics Consortium (TAG) (20), we obtain some interesting results.

## MATERIAL AND METHODS

### QCAT assuming single causal variant

The proposed approach uses a fast regression procedure to assess the likelihood of any cataloged (regardless whether GWAS measured or unmeasured) SNP being causal. It relies solely on the association statistic at the SNP, the statistics for the adjacent SNPs at/near the loci and the genotype correlation matrix between all these SNPs, e.g. as estimated from a reference panel such as the 1KG data.

Let *G_c_* and *Z_c_* be the genotype vector and the association Z-score for **measured/unmeasured** putatively causal SNP under investigation, *c* = 1, …, *N*, where *N* is the number of SNPs in a region of interest. Let *G_i_* and *Z_i_ i* = 1, …, *M*, be the genotype vectors and association Z-scores for all **measured** SNPs in an extended window around SNP *c*. Let *ρ*_*ci*_, *i* = 1, …, *M*, be the correlation between *G_c_* and *G_i_* estimated from the reference population. Under/near the null hypothesis (H_0_) of no association between SNPs and disease and, when **testing an additive mode of inheritance** (MOI), **ρ*_ci_* is also the correlation between *Z_c_* and *Z_i_*. If the SNP *c* is causal, then for every measured non-causal SNPs in the extended window, we can write E(*Z_i_*|*Z_c_*) = ρ_ci_ *Z_c_* or, in regression notation, *Z_i_* = ρ_ci_ *Z_c_* + ε_*i*_ (1), where ε_*i*_, *i* = 1, …, *M* are the non-independent (and non-identically distributed) errors due to *Z_i_* being locally correlated. Relationship (1) implies that, if is causal with the assumed MOI, then the correlation (regression) between vectors **Z** = {*Z_i_*} and ***ρ***_c_ = {*ρ_ci_*}, *i* = 1, …, *M*, should be non-zero, i.e. *cor*(***Z, *ρ*_c_***) ≠ **0**. This test can be viewed as a **test for an additive MOI association under the assumption of causality**. Thus, in practice we can test for (univariate) causality all SNPs in large reference panels (e.g. 1KG) by testing *cor*(***Z, *ρ*_c_***) ≠ **0**, henceforth denoted as QCAT.

Under H_0_ (no significant association under the assumption of causality): *cor*(***Z, *ρ*_c_***) = **0**, the empirical distribution of the sample correlation between the dependent variable, ***Z***, and LD parameters, ***ρ***_*c*_, is known only when entries of vector are independent, which is not the case when the SNPs are in LD. However, given that the correlation matrix (LD) between SNP genotypes/Z-scores (Σ) can be estimated from the reference population, the observations can be transformed to independence by left multiplying ***Z*** and ***ρ_c_*** with *Σ*^−1/2^ (i.e. Cholesky transformation. Under a full rank transformation, Invariance Principle of MLE (21) surmises that testing *cor*(***Z, *ρ*_c_***) = 0 is equivalent to testing 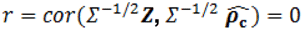. To assess its significance, we can use the correlation (regression) test of transformed data based on 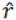 (the sample estimate of *r*):

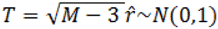 or 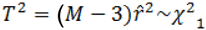.

The rest of the paper deals chiefly with the above described single causal variant QCAT. In this form, QCAT should cover a large number of practical scenarios. However, its extension to multiple causal SNP scenario is straightforward, as described in the first section of the Supplementary Material (SM).

### Applying QCAT to whole genome

In addition to testing for causality SNPs in a particular genomic region, QCAT software also offers sequential QCAT procedure automatically testing variants through the whole human genome by using a sliding window strategy, as adopted in our other summary statistic based tools (22;23). For each iteration, QCAT determines non-overlapping test window with a fixed size (1 Kb by default) and extended window including the test window along with fixed-size (150 Kb by default) upstream and downstream regions. Then, QCAT tests for causality SNPs within the test window using Z-scores of SNPs in the extended window and LD matrix estimated from homologous reference SNP genotypes.

### Extension to cosmopolitan cohorts

QCAT performs best when the LD structures of both study cohort and external reference population are identical. To make it applicable to summary data from mixed ethnicity cohorts, similar to our DISTMIX and JEPEGMIX tools (18;19), we estimate the cohort LD (*Σ*) as the mixture of the LDs of the reference panel ethnicities using user-provided/estimated ethnicity proportions. In this paper we will refer to such version as QCAT for MIXed ethnicity cohorts (QCATMIX).

### Type I error rates

We assessed the Type I error rates of the proposed method for both homogeneous and mixed ethnicity cohorts under H0 of no association between phenotype and SNP genotypes. For each scenario we performed 100 simulations of 10,000 individuals which were sampled with replacement from 1KG panel (24) i) European cohort for the homogeneous scenario and ii) all subjects for the mixed ethnicity scenario. Genotype data for each such cohort was analyzed only for SNPs with minor allele frequency >1% in a randomly selected autosomal region with a length of 50 Kb. We simulated the phenotype data using a random standard normal distribution. Summary statistics (two-sided Z-scores) were obtained by simply regressing the simulated SNP genotypes on the phenotypes. Simulated data sets were analyzed using both QCATMIX and the commonly used regression association method. Since CAVIAR, CAVIARBF and PAINTOR are designed for posterior probability prediction and not statistical testing, we are not able to assess their size of the test via such simulations.

### Comparison with other fine-mapping methods

For various patterns and strength of LD, we performed 1,000 simulations to compare the performance in fine mapping of QCAT(MIX), CAVIARBF (C-BF), as a representative of posterior probabilities methods, and GCTA-COJO (denoted simply as COJO). While COJO does not really infer causality, it is used as an illustration of the conditional approach. For each simulation, we simulate Z-scores for 100 SNPs assuming that one of the two in the middle, denoted as k, is the causal marker, which has a Z-score of μ = {0, 2.5, 5, 10, 15, 20, 25}. Given that this SNP was assumed causal, the Z-score for the *i*-th SNP is obtained simply as *Z_i_* = *ρ*_*ik*_ μ, where *ρ*_*ik*_ is the nuisance correlation between the genotypes for the two SNPs. We simulated four patterns of LD between the 100 Z-scores: i) autoregressive (AR), *ρ*_*ij*_ = *ρ*^|*i−j*|^, ii) exchangeable (EX) *ρ*_*ij*_ = *ρ*, iii) half-AR, 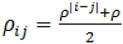, and iv) block AR, i.e. SNPs form 10 block of 10 SNPs and their correlations are EX within block and AR between blocks. As strength of the LD in the above LD patterns we investigated *ρ* = {0.8, 0.9, 0.95, 0.99}. Besides analyzing all simulated SNP set, to simulate the uncatalogued causal variants, we also analyzed just non-causal SNPs. Under the alternative hypothesis experiment, we also compute the LD between QCAT Z-scores and contrast them to association.

### Practical application

For a more empirical performance assessment, we used summary data sets for smoking quantity (number of cigarettes smoked per day) from TAG study (TAG CPD) (20). This study yielded the well known, large signal in nicotinic receptor gene cluster (15q25.1). For QCAT we thus used summary statistics for SNPs from this region with a window size of 250Kbp.

## RESULTS

As C-BF and related methods are all just tools for predicting the posterior probabilities of causality (under simplifying assumptions), type I error rates are applicable only to QCAT and association methods. Similar to our previous methods, e.g. DIST/DISTMIX and JEPEG/JEPEGMIX, QCAT is somewhat conservative (Fig. 1). However, we expect its conservativeness to diminish with an increase in reference population panel size.

**Figure 1.**
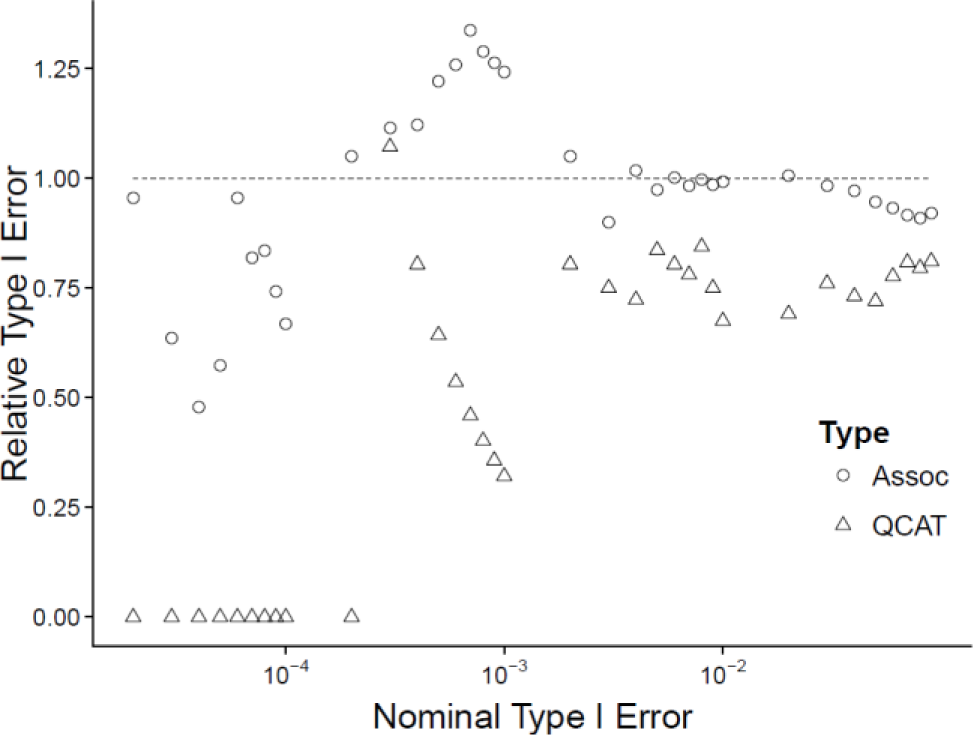
Type I error rate for association (circle) and QCAT (triangle).

The alternative hypothesis simulations estimated the probability of detecting any signal at “genome-wide” significant levels, i.e. for QCAT and COJO at a genome-wide type I error of 5×10^−8^ and, for C-BF, at a probability of a causal variant (in the set of 100 SNPs) >1- 100×5 10^−8^. The only exception to the above rule is for the scenario when the causal SNP is measured, in which QCAT conservatively tests only the causal marker. Simulations show (Fig. 2) that when the causal SNP is not measured (Causal=OUT), the probability of detecting at least one causal SNP is rather high for both CAVIARBF and COJO (Fig. 2 dashed lines), often even for a genome-wide nonsignificant signal at the causal SNP (μ = 5). The probability of detection for C-BF and COJO is little changed between the scenarios of the causal SNP being in (Fig 2 solid lines) or out. However, for QCAT, the probability of detecting the exact causal SNP increases when it is measured (solid lines) as opposed to detecting at least one causal marker in the region when the causal SNP is not measured (dashed lines). The increase is substantial for LD structures other than AR.

**Figure 2.**
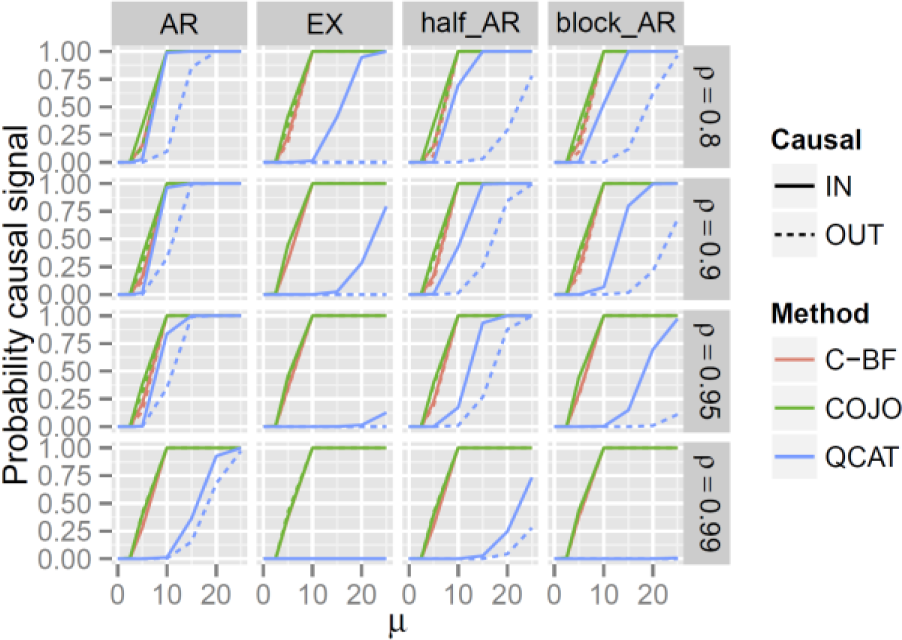
Probability of detecting a causal signal for various methods when the true causal SNP is measured (solid line) and unmeasured (dashed line).

The LD of QCAT statistics and neighboring SNPs is very different from the LD of association statistics (Fig. 3). The LD of QCAT statistics decays much faster. The decay rate contrast is especially notable for EX models and/or larger levels of LD between neighboring markers (*ρ*).

**Figure 3.**
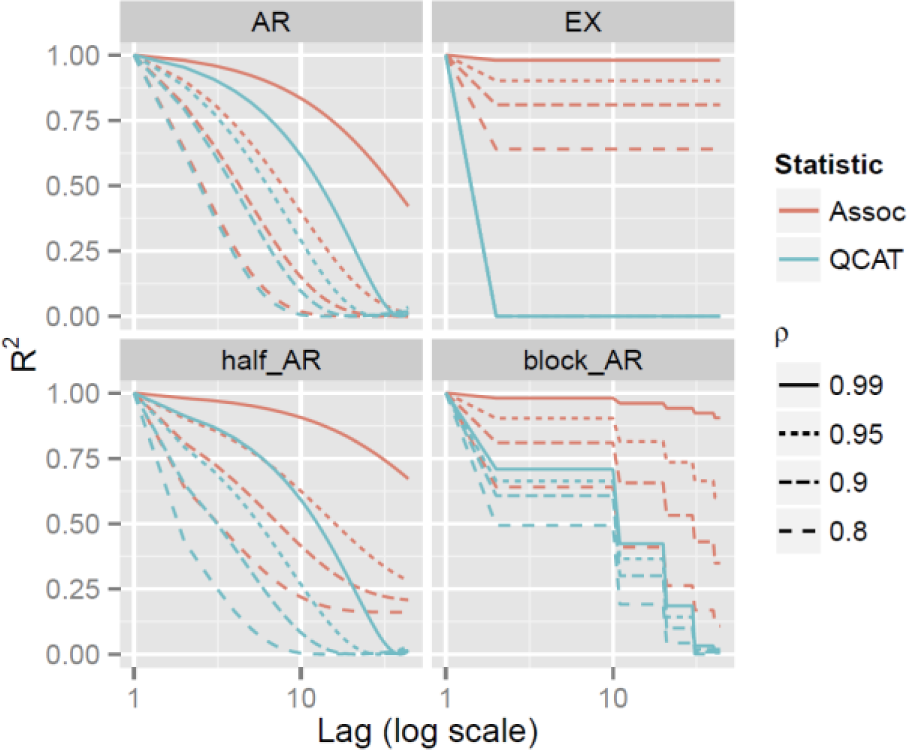
LD for association and QCAT statistics (from 100 SNP window) for neighboring SNPs.

**Figure 4.**
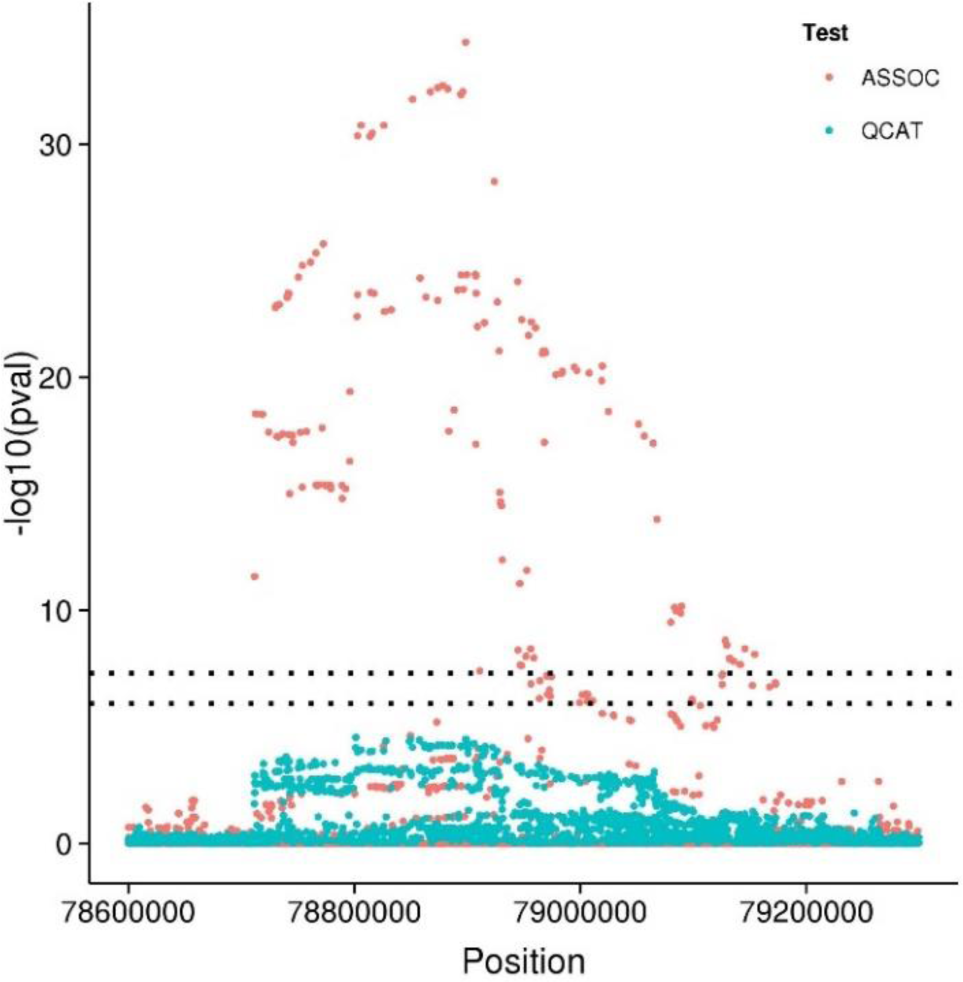
TAG association (red) and QCAT (green) pvalues by position (bp) for the nicotinic receptor gene cluster (15q25.1).

The practical application to SNPs presented in the TAG study (Fig. 3) shows that while the best association signal in this GWAS is strong, and tools like CAVIARBF predicting it as being almost surely causal, the QCAT signal is not significant (albeit reasonably close). While further investigations are necessary and underway, this leads to a very nuanced initial suggestion that either i) the causal variant is not measured in the study, i.e. researchers need to look among non-GWAS/non-imputed variants, or ii) the MOI of the causal variant is not additive.

## DISCUSSION

We propose a novel tool for fine-mapping which, unlike to existing methods, i) statistically tests variants (as opposed to just predicting the posterior probability) for causality and ii) does not need to assume that the causal variants are measured/catalogued. The test is based on the observation that a causal variant with a significant signal induces a correlation between Z-scores of measured SNPs and their correlation to the causal variant. The method maintains type I error at or below nominal levels.

To avoid spurious results, similar to our other software, DISTMIX and JEPEGMIX, the correlation matrix is computed from the reference panel using a ridge estimate. This results in QCAT being rather conservative. However, the ridge penalty is sample size dependent and, with an expected substantial increase in sample size for future reference panels, it will become much smaller. Thus, with such reference panels, QCAT is likely to be only slightly conservative.

While QCAT, due to its testing approach, shows better performance than other competitors, simulations under alternative underscore the many possible pitfalls of statistical fine-mapping procedures. At large LD levels between SNPs, even QCAT cannot detect the difference between causal and non-causal SNPs. Thus, fine-mapping methods seem to be most useful only when applied to SNPs/components in moderate LD. To achieve this in practical applications of dense SNP data might be achieved by a four step process. First, cluster SNP at a moderate level (e.g. *r*^2^ = 0.5 − 0.6). Second, take as representative for the cluster the first principal component of SNPs in the cluster, possibly using weights dependent on functional information. Third, to find clusters harboring causal mutations, apply fine-mapping methods like QCAT to cluster representatives. Fourth, in-vivo validation means, e.g. apply CRISP(R) mutations in cell cultures, to determine which variant in the cluster is causal.

In its present form, QCAT performs only the univariate test for causality and does not use SNP functional information. Conceptually, its extension to multiple variants should be simple (see SM). Similarly, it should reasonably straightforward to extend the method to use pleiotropy to enhance the test, e.g. along the lines of Pickrell et al (25). However, a through inclusion of functional information might be substantially more involved. Until such approach is available, researchers might compute variant weights based on functional information and use them, along with QCAT p-values, in a weighted FDR procedure.

Similar to competing tools, the current QCAT version has the main limitation of *assuming an additive MOI*. Thus, a non-significant test of causality can be, as seen from the practical application, due to the variant not being causal *or* the MOI being far from additive. Therefore, great care should be taken in interpreting QCAT results. However, in future versions of QCAT we plan to test more general MOIs. This might be achieved by a data dependent estimation of where the best fitting model sits in the continuum between the dominant and recessive (even over-dominant) MOIs.

Due to the LD of QCAT tests decaying much faster than their association counterpart, QCAT statistics are prime candidates for constructing multi-variant statistics. Interesting instances of such statistics might be enrichment type tests such as gene-, pathway-, tissue mark-level etc. However, given that the LD pattern of QCAT statistics is rather complex, constructing multivariate statistics could be rather laborious.

## FUNDING

This work was supported by the National Institutes of Health [DA026119 to D.L., MH100560 to B.P.R. and S.A.B., AA022717 to V.I.V. and S.A.B]. Funding for open access charge: National Institute on Alcohol Abuse and Alcoholism.

## SOFTWARE

The QCAT software is publically available at: http://dleelab.github.io/qcat/.

## Supplementary Material

### 1 Extension to multiple causal variants

While the paper deals with single causal SNPs (which should covers a large number of practical scenarios), QCAT can be easily extended to multiple, reasonably close causal variants. (In practice these causal SNPs might be detected heuristically, for instance, one-at-a-time by applying the single SNP QCAT.) Thus, when there are interesting signals for SNPs that are relatively close, testing all these SNPs simultaneously might be useful. To derive a multi SNP QCAT, with the notation from the previous section, let the *k* SNPs, *c*_1_, …, *c*_*k*_, be the indices for causal SNPs. Then we can write 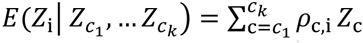 (2). I.e., we can test whether *Σ*^−1/2^***Z*** is independent from any linear combination of 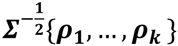. This is the ubiquitous test of all slopes being zero in the simple multiple regression of *Σ*^−1/2^***Z*** on 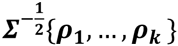 (Anderson, 2003). The test for zero slopes is asymptotically distributed as a χ^2^ with *k* df (*i.e.* χ^2^_*k*_).

The regression approach for multiple variant QCAT test also opens the possibility of treating the prediction of causal variants as a simple model selection for classical linear models. For instance, a forward model selection, similar to step.lm from R, within a sliding window can be employed for predicting the likely causal SNPs. Given that the number of causal SNPs in a window is likely to be small, such an approach will be very computationally efficient by not testing sets with a large number of SNPs.

